# SPAED: Harnessing AlphaFold Output for Accurate Segmentation of Phage Endolysin Domains

**DOI:** 10.1101/2025.04.25.650745

**Authors:** Alexandre Boulay, Emma Cremelie, Clovis Galiez, Yves Briers, Elsa Rousseau, Roberto Vázquez

**Affiliations:** Department of Biotechnology, Ghent University, Ghent, Belgium; Département de biochimie, de microbiologie et de bio-informatique, Université Laval, Québec, QC, Canada; Centre Nutrition, Santé et Société (NUTRISS), Institute of Nutrition and Functional Foods (INAF), Université Laval, Québec, QC, Canada; Univ. Grenoble Alpes, CNRS, Grenoble INP, LJK, 38000 Grenoble, France; Département d’informatique et de génie logiciel, Université Laval, Québec, QC, Canada; Centre de Recherche en Données Massives de l’Université Laval, Québec, QC, Canada; Institut Intelligence et Données (IID), Université Laval, Québec, QC, Canada; Centro de Investigación Biomédica en Red de Enfermedades Respiratorias (CIBERES), Madrid, Spain

## Abstract

**Summary:** SPAED is an accessible tool for the accurate segmentation of protein domains that applies hierarchical clustering to the predicted aligned error (PAE) matrix obtained from AlphaFold predictions. It leverages information contained in the PAE matrix to better identify domain-linker boundaries and detect disordered regions. On a dataset of 376 bacteriophage endolysins (proteins that degrade the bacterial cell wall), SPAED achieves a mean intersect-over-union score of 96% and a domain-boundary-distance score of 89% compared to 94% and 70%, respectively, for the state-of-the-art tool Chainsaw.

**Availability and Implementation:** SPAED is available on the web at http://spaed.ca and available for download at https://github.com/Rousseau-Team/spaed.

**Contact:** Elsa Rousseau -, Roberto Vázquez -

## Introduction

Bacteriophages (phages), viruses that infect bacteria, are the most abundant biological entities on Earth^1^. The ongoing coevolution between them and their bacterial counterparts has led to a vast diversity of phages that are in constant evolution^2^. To release their progeny in the environment, phages often rely on endolysins that degrade the peptidoglycan layer of the bacterial cell wall thus enabling lysis of the host^3,4^. The complexity and variety of bacterial cell wall architectures – a result of the diverse peptidoglycan, teichoic acids, lipopolysaccharides and different substituents composing it – have driven phages to refine their lytic cassette to be tailored to their host^5^. Consequently, the diversity of endolysins mirrors that of the phages themselves and of their hosts^6,7^.

Many phage endolysins have a modular structure, with each module usually possessing either a cell wall-binding or catalytic function^6,8^. In nature, this modularity enables phages to evolve their endolysin gene, by acquiring new domains by recombination^9^. Similarly, novel lysins can also be engineered in a lab by domain shuffling to enhance lytic activity, binding affinity, specificity, etc.^5,8,10^. Thus, the accurate identification of lysin domains is important from a biological perspective as well as for the development of new antimicrobial agents^8,11^.

Existing tools for protein domain segmentation are not specifically adapted to endolysins. Since the advent of AlphaFold2/3^12,13^, many state-of-the-art tools use supervised deep learning models trained on large datasets and based on structural information^14–16^. They are shown to work well in general but depend on the quality of annotations present in these databases. Although they have grown in recent years, protein domain databases are not necessarily representative of all modular proteins, and phage proteins are particularly underrepresented^17,18^. In contrast to supervised approaches, unsupervised-heuristic algorithms have also been used historically, sometimes in combination with information from annotated domain databases^19,20^, but these approaches typically struggle to encompass all cases.

Here we developed SPAED, a tool for the **S**egmentation of **P**h**A**ge **E**ndolysin **D**omains that applies hierarchical clustering to the predicted aligned error (**PAE**) matrix obtained from AlphaFold predictions. The PAE is a score that estimates the expected positional error for each pair of residues in a predicted protein structure by calculating the error associated with aligning each residue to every other^21^. It is a measure of the local packing of residues in a protein as well as of the relative placement of domains in the predicted structure. SPAED uses these expected positional errors as a measure of how likely residues are to be found in the same domain. This approach is well suited for the identification of endolysin domains because they are mostly compact and separate from one another, which is reflected in the PAE matrix of these proteins. SPAED was tested extensively on a dataset of 376 manually delineated endolysins and we also demonstrate its applicability to other types of modular proteins obtained from CASP12^22^. SPAED can easily be launched from our web portal available at www.spaed.ca and is downloadable through GitHub and PyPI for ease of use on larger datasets.

## Methods

A dataset of 376 endolysins, visually delineated based on predicted 3D structures, was obtained from previous and ongoing projects performed at Ghent University^7,23,24^. The 3D structures of all lysins were predicted using ColabFold v1.5.5^25^ and the PAE files were collected. The whole dataset is available as supplementary material.

### Algorithm

A complete example of the algorithm with explanations and visuals for each step is shown in Supplementary material Appendix 1.

At the basis of SPAED is a single linkage hierarchical clustering algorithm (hierarchy.fclusterdata from SciPy) that takes as input a symmetrized PAE matrix from AlphaFold (Figure 1A; step 1). This symmetrized matrix is obtained by averaging the PAE matrix and its transpose (pae + *pae*^*T*^)/2. Applying the clustering to the columns of the resulting matrix places residues with similar profiles in the PAE matrix into the same cluster. In practice, there is a certain degree of noise in the PAE matrix that requires adjustments to sensitively detect domain boundaries.

**Figure 1.**
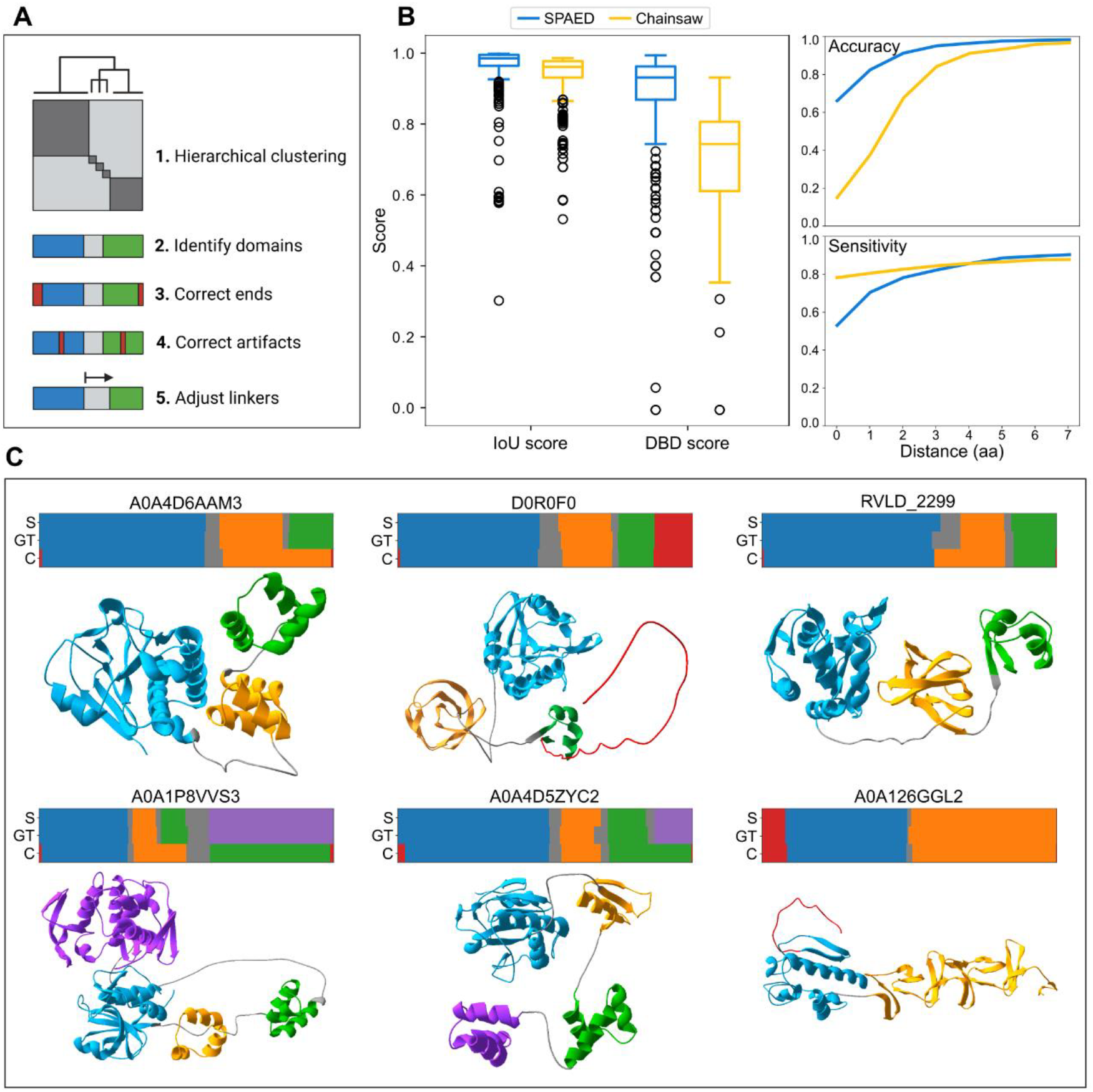
**A)** SPAED algorithm overview. **B)** SPAED and Chainsaw’s performance on the 376 lysin dataset. IoU: Intersect over union. DBD: Domain boundary distance. Accuracy and sensitivity are shown as a function of the distance (in amino acids, aa) between the predicted and ground truth (GT) boundaries. **C)** Comparison of SPAED (S), GT and Chainsaw (C) domain segmentations for 6 endolysins representative of different architectures. 3D structures show domains predicted by SPAED and were made using SwissPdb Viewer^26^. Disordered regions are in red, linkers are in grey, and domains are represented in other colors (blue, orange, green and purple).

The hierarchical clustering is restricted to a maximum number of clusters (criterion = “maxclust”) that we set to 1/10^th^ of the length of the protein after a series of tests (see performance_eval.ipynb in the GitHub repository), typically resulting in 20 to 60 clusters. This number is high compared to the expected number of domains as lysins are known to possess 1-4 domains. However, this allows a necessary flexibility to the clustering algorithm which then assigns a high number of small, preliminary clusters in less compact regions, such as the extremities of the protein and between domains (i.e., linker regions). As a result, many small, often-singleton clusters are produced in those regions, whereas long, structured clusters are generated in more organised regions.

Clusters containing more than 25 residues are assigned as predicted domains (Figure 1A; step 2). This number accounts for the smallest expected size of a domain (>30 residues)^27^ and leaves a buffer for errors in the preliminary assignment of clusters. All other clusters (containing <25 residues) are assigned a “non-domain” identifier. According to their position, these “non-domain” residues can be **1)** terminal disordered regions when they are found at either end of the protein, **2)** wrong assignments (artifacts) when they are found within a single domain or **3)** linkers when they are found between two domains. These non-domain regions are dealt with separately within the algorithm:

1. Terminal regions labelled as “non-domain” are concatenated to the nearest domain if they are less than 20 residues long (Figure 1A; step 3). Alternatively, an additional filter is applied to evaluate if the region is truly disordered, in which case it could correspond to a signal peptide of interest (usually at least 25 residues long)^28,29^. Such a filter works by counting the number of neighbours of each residue in the region with a “low” PAE score (< 5). The latter threshold was defined through experimental observation on the lysin dataset (see Supplementary figure A1.3) and has been used before as a measure in other tasks^30^. Then, if >80% of residues in the region have 5 neighbours or less with low scores, then the region is considered disordered. If the region is deemed to be ordered, it is concatenated to the nearest domain as it likely corresponds to a less packed (but still ordered) region of that domain.
2. Artifacts, or simply small errors that originate from the original clustering, are then removed (Figure 1A; step 4). For this, a simple sliding window is used to verify that all domains are continuous, making the necessary adjustments if that is not the case.
3. Finally, the boundaries of the linkers are adjusted (Figure 1A; step 5). Similarly to how terminal disordered regions are detected, residues that are part of a linker will have a low PAE score (PAE < 5) with less residues than those that are a part of a domain. Conversely, residues found in a domain should have a low PAE score with about as many residues as are part of that domain. Thus, residues near the domain-linker boundary are considered part of the domain if they have a low PAE score with at least 25 residues (a domain/repeat is expected to be >30 residues long)^27^.

### Evaluating and comparing performance

To validate results on our dataset of 376 lysins, we compared the predictions made by SPAED to those made by the manual annotation (ground truth; GT) and to those made by the most recent, state-of-the-art model for protein domain segmentation called Chainsaw^14^. Chainsaw is a structure-based, supervised method that employs a convolutional neural network (CNN) to estimate the probability that pairs of residues belong to the same domain. It was shown to outperform other segmentation tools such as Merizo^16^, EguchiCNN^15^, UniDoc^31^ and SWORD2^32^.

We first compared the quality of segmentations using the intersect over union (IoU) score (Supplementary material Appendix 2)^33^. This score is a measure of the overlap between GT and predicted domains (with a score of 1 corresponding to a perfect overlap)^14^. We also evaluated the accuracy of the predicted boundaries using the Domain Boundary Distance (DBD) score^34^. This score rewards a predicted boundary that is closer to the GT boundary; one point is attributed for a perfect prediction and 1/8 point is subtracted for every residue between the predicted and GT boundary.

## Results and discussion

We benchmarked SPAED against Chainsaw^14^ on a dataset of 376 endolysins by comparing the predictions made by both tools to the manual annotations made by an expert and based on the predicted 3D structures. The IoU and DBD were used as evaluation metrics. SPAED averages an IoU-score of 96% ± 8% (SD) compared to 94% ± 7% for Chainsaw (Figure 1B). A larger difference is observed with the DBD-score where SPAED achieves an average score of 89% ± 15% and Chainsaw has a score of 70% ± 14%.

The DBD-score measures both the accuracy (fraction of predicted boundaries that are correct) and sensitivity (fraction of true boundaries that are correctly predicted) of predictions^34^. Viewing both separately is also essential to avoid emphasizing completeness at the expense of accuracy (over-predicting boundaries) or sacrificing completeness in favour of accuracy (under-predicting boundaries). Figure 1B shows the accuracy and sensitivity of boundary predictions, calculated over a permissibility range that allows for distances of 0 to 7 residues between the predicted and true boundary when classifying a prediction as correct/incorrect. More than 65% of domain boundaries are predicted exactly by SPAED (at distance between predicted and GT = 0) and, given a buffer of 2-4 residues, nearly all boundaries are predicted correctly. A remarkable (50%) drop in accuracy is observed when comparing Chainsaw to SPAED for exact predictions (Figure 1B at distance = 0), but an important improvement is noticed when tolerating predictions up to 7 residues off. Regarding sensitivity, Chainsaw is better at a distance of 0 to 3 residues, but when allowing a 4-5 residue buffer, SPAED becomes marginally better using this metric as well. Given that various parameters (PAE score, maximum number of clusters, etc.) were optimized for SPAED on the same endolysin dataset, a slight bias in the reported metrics may be observed for our tool, however, these parameters do not drastically affect SPAED’s performance overall (see performance_eval.ipynb in the GitHub repository).

The accuracy of SPAED predictions can be seen in Figure 1C for 6 endolysins representative of different domain architectures. The examples also highlight some minor flaws in Chainsaw’s predictions, such as repeated domains that are often ignored by Chainsaw (A0A4D6AAM3, A0A1P8VVS3, A0A4D5ZYC2). In addition, SPAED can identify disordered regions, potentially signal peptides, in N- or C-termini (Figure 1C, colored in red). These can sometimes be recovered from Chainsaw predictions if the regions were not assigned to any domain, but they are generally ignored by the tool. In addition, SPAED parameters can be adjusted to detect disordered regions and linkers more sensitively if needed.

Although SPAED was built for endolysins, it can also be used on other types of proteins. As a test, 18 modular proteins were collected from the CASP12 experiment^22,35^. Their 3D structures and domains were predicted using AlphaFold3 and SPAED, respectively. These, as well as the GT delineations obtained from CASP12 when available, can be visualized in Supplementary Figure A3. Note that SPAED parameters are tunable and were adjusted for some of these proteins as specified in the figure. For 12 proteins (A-F, L-Q), the delineations obtained by SPAED are accurate. Errors in the remaining 6 proteins result from tightly packed domains (H, J, K) or discontinuous domains (G, I, R) that complicate the detection of boundaries in the PAE matrix. SPAED was also applied to two cellulosome components (a docking enzyme and a scaffoldin, Supplementary Figure A3 panels S, T) and good delineations were obtained^36^. Like endolysins, their domains are compact and separate from one another, making SPAED well-suited for their accurate delineation.

To conclude, SPAED allows for the high-throughput segmentation of protein domains in a simple and interpretable manner. It is also flexible in that its parameters can be modified to more sensitively detect linkers or disordered regions, or to improve segmentations if needed (e.g., for lower throughput experiments). Additionally, users can submit a folder of PAE files to annotate multiple proteins simultaneously, and it is possible to get a 3D visualisation of the predicted domains by adding the protein structure files on a user-friendly website (www.spaed.ca), making it accessible to users less familiar with bioinformatics. Although it was initially developed for and optimized on endolysins, SPAED can be used on other types of modular proteins characterized by compact and relatively distant domains.

## Acknowledgements

AB was supported by fellowships from the FRQNT (#325947), from the CREATE Responsible Health and Healthcare Data Science (RHHDS) program from NSERC and from the Mitacs Globalink Research program. ER was funded by a Research Scholars – Junior 1 in artificial intelligence and digital health by FRQS (#307935). RV was supported by a postdoctoral fellowship of the ‘Bijzonder Onderzoeksfonds’ (BOF), Ghent University (01P10022). EC was funded by Research Foundation – Flanders (FWO) (1S15424N). This research was enabled in part by support provided by Compute Ontario (https://www.computeontario.ca/) and the Digital Research Alliance of Canada (alliancecan.ca).

Figure 1A was created in BioRender.

## Conflict of interest

YB is co-founder and scientific advisor of Obulytix. RV has provided scientific consulting services to Obulytix.

## Contributions

**AB**: conceptualization (equal); software (lead); writing – original draft (lead); formal analysis (lead); writing – review and editing (equal). **RV**: conceptualization (equal); data curation (lead); writing – review and editing (equal); supervision (lead). **ER**: writing – review and editing (equal); supervision (equal). **YB**: writing – review and editing (equal); supervision (equal). **CG**: software (supporting); writing – review and editing (equal). **EC**: conceptualization (supporting); software (supporting); writing – review and editing (equal).

## Notes

https://doi.org/10.5281/zenodo.15285861

### Competing Interest Statement

Yves Briers is co-founder and scientific advisor of Obulytix. Roberto Vazquez has provided scientific consulting services to Obulytix.

