## Supplementary material for "SPAED: Harnessing AlphaFold Output for Accurate Segmentation of Phage Endolysin Domains"

### Appendix 1: Complete example of the SPAED algorithm.

Here is the example endolysin we will be working with.

**Supplementary figure A1.1**:

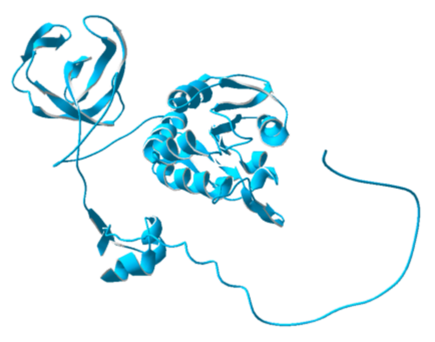

#### Preprocessing

SPAED takes as input the **PAE matrix** obtained from AlphaFold.

The matrix is normalized to make it symmetric:

$$pae_{norm} = \frac{(pae + pae^T)}{2}$$

**Supplementary figure A1.2**:

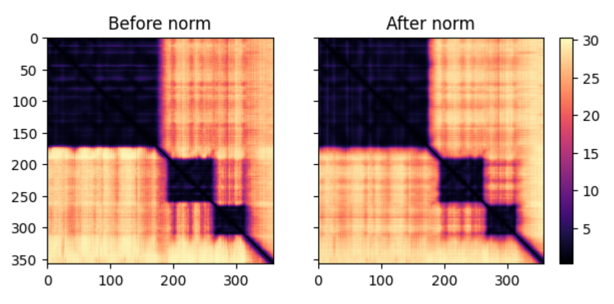


#### Step 1: Hierarchical clustering

Hierarchical clustering (scipy.cluster.hierarchy.fclusterdata) is performed on the normalized matrix. The maximum number of clusters is set to 1/10$^{th}$ of the length of the protein, so in this case to 35 clusters. This step places the most similar columns of the PAE matrix into the same clusters.

The article refers to the **profile** of a residue in the PAE matrix. The profile of a residue is obtained by taking all the values in its corresponding column of the PAE matrix. Residues in the same domain should have similar profiles. Also, notice that within a domain, residues mostly have a PAE score <=5.

**Supplementary figure A1.3**:

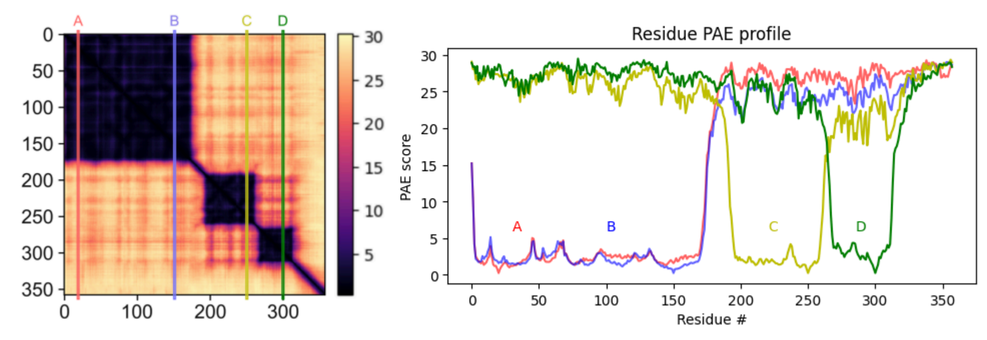


Applying the clustering algorithm yields the following result, where each colour corresponds to a different assigned cluster.

**Supplementary figure A1.4**:

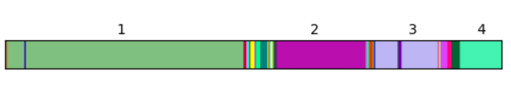


At first glance, we can already see 4 clusters, each containing many residues.

Notice how residues in linker regions are assigned to many small clusters. This is a result of the high number of clusters (in this case 35) that the clustering algorithm is allowed to look for. In contrast, long, continuous clusters are assigned to more homogeneous regions of the PAE matrix.

#### Step 2: Domain detection

Clusters with 25 or more residues are assigned as domains. This threshold was chosen as domains are expected to have at least 30 residues. A little buffer is given for errors present in the preliminary assignment of clusters.

Each identified domain is assigned a new cluster number (1-4 in this case) whereas all other residues are assigned a 'non-domain' id.

**Supplementary figure A1.5**:

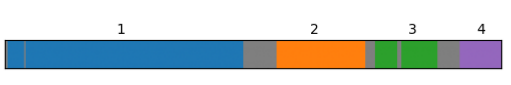


#### Step 3: N- and C-terminal correction

Next, we correct the ends of the sequences. These regions often possess residues that are freer to move and thus have a profile in the PAE matrix that is very different from residues in the domain they are nearest to. If this region is very short (<20 residues), it is simply concatenated to the nearest domain (like the first residue in this example).

Endolysins can also possess signal peptides in N- or C-terminal which correspond to longer (~25 residues) disordered regions. These are interesting to flag for future studies. In the PAE matrix, these disordered regions can be identified by looking for low PAE score 'diagonals'. In practice, we can count the number of 'dark' squares (PAE score <5) in every column (see figure 3 for the reason for using PAE score <5). This gives a good estimation of the packing of the protein in any given region as it reflects the number of residues that are close in space to any given residue. More importantly, residues in less packed regions such as linkers or end-terminal disordered regions will have very few of these 'dark' squares (i.e. they correspond to the 'diagonal' regions).

**Supplementary figure A1.6**:

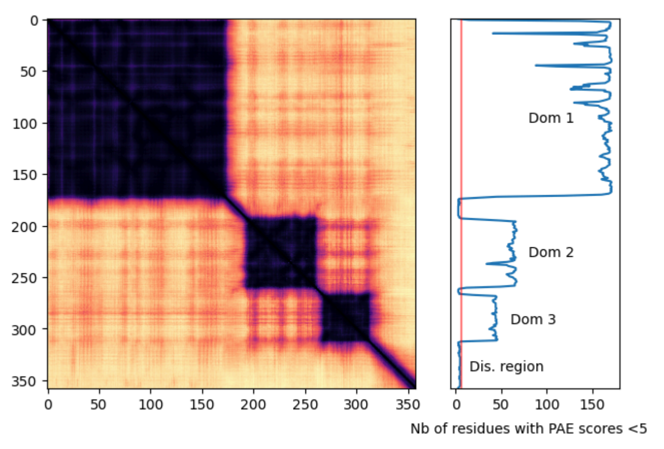


If more than 80% of residues in a region are below the cutoff (red line in the graph above) corresponding to 6 residues with scores < 5, the region is considered to be disordered.

The new assignment yields the following, where the 4th region, now in red, is flagged as being disordered.

**Supplementary figure A1.7**:

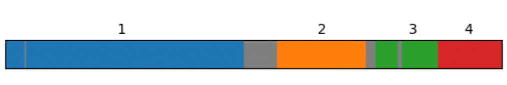


#### Step 4: Artifact removal

We correct small artifacts originating from the initial assignment of clusters by scanning for residues assigned as 'non-domain' within a domain. This fixes the wrongly assigned residues like in domain 3.

**Supplementary figure A1.8**:

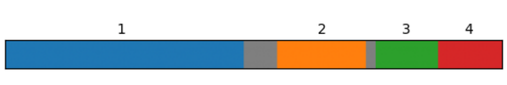


#### Step 5: Linker adjustment

Finally, we adjust the length of linkers. This step is necessary to correct small mistakes at the domain/linker interface that can originate from the initial clustering.

Residues near the domain/linker boundary are considered part of the domain if they have a low PAE score (<5) with at least 25 residues (a domain is expected to be at least 30 residues long) as residues found in a domain should have a low PAE score with about as many residues as are part of that domain.

**Supplementary figure A1.9**:

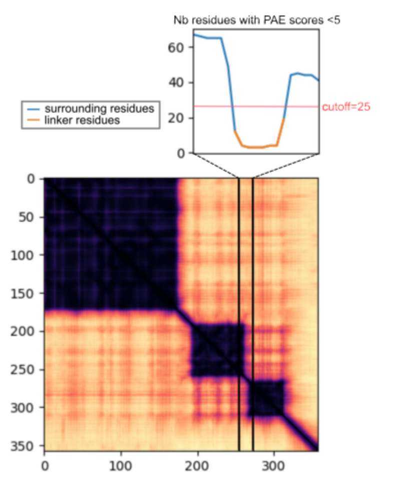


**Supplementary figure A1.10**:



#### Final result

Finally, we can see how the predicted domains look on the predicted 3D structure.

**Supplementary figure A1.11**:

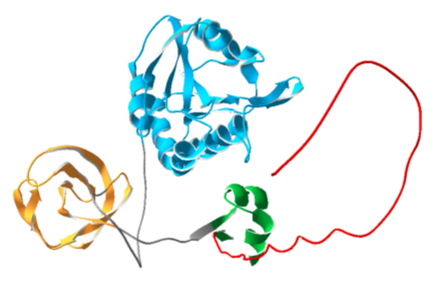

### Appendix 2: Extra explanation of Intersect over Union (IoU) score.

As described in Wells et al.[14]: following the approach of Merizo[16], for a given protein chain we compute the average intersection over union (IoU) between paired sets of predicted and ground-truth residues assigned to each domain. As a first step, each ground-truth domain is paired with a predicted domain such that the sum of all intersections over unions is maximised while respecting the following constraints: Each ground-truth domain (represented as a set of residue indices) Ti can have, at most, one paired predicted domain Pi . Second, each predicted domain can be assigned at most once. No IoU is computed for the sets of residues that are labelled as, or predicted to be non-domain residues. To generate a final score for the whole chain each domain-level IoU is weighted by the number of residues in the ground-truth domain:

$$IoU = \sum_{i=1}^{n_{dom}} \frac{|T_i \bigcap P_i|}{|T_i \bigcup P_i|} \cdot \frac{|T_i|}{\sum_{j=1}^{n_{dom}} |T_j|}.$$

### Appendix 3: Example segmentations of CASP12 proteins and cellulosomes.

**Supplementary figure A3**: Comparison between domain delineations obtained with SPAED and the ground truth (GT) for proteins from the CASP12 experiment (**A-R**), as well as two example cellulosome components (**S, T**). SPAED was launched with default parameters unless specified otherwise. Segments in red correspond to disordered regions. Segments in grey correspond to linkers, except in **A** where a large segment was undefined in the GT. The GT delineations could not be retrieved for **L-R** but all structures are described as having 2 domains in Zhou et al.[35].

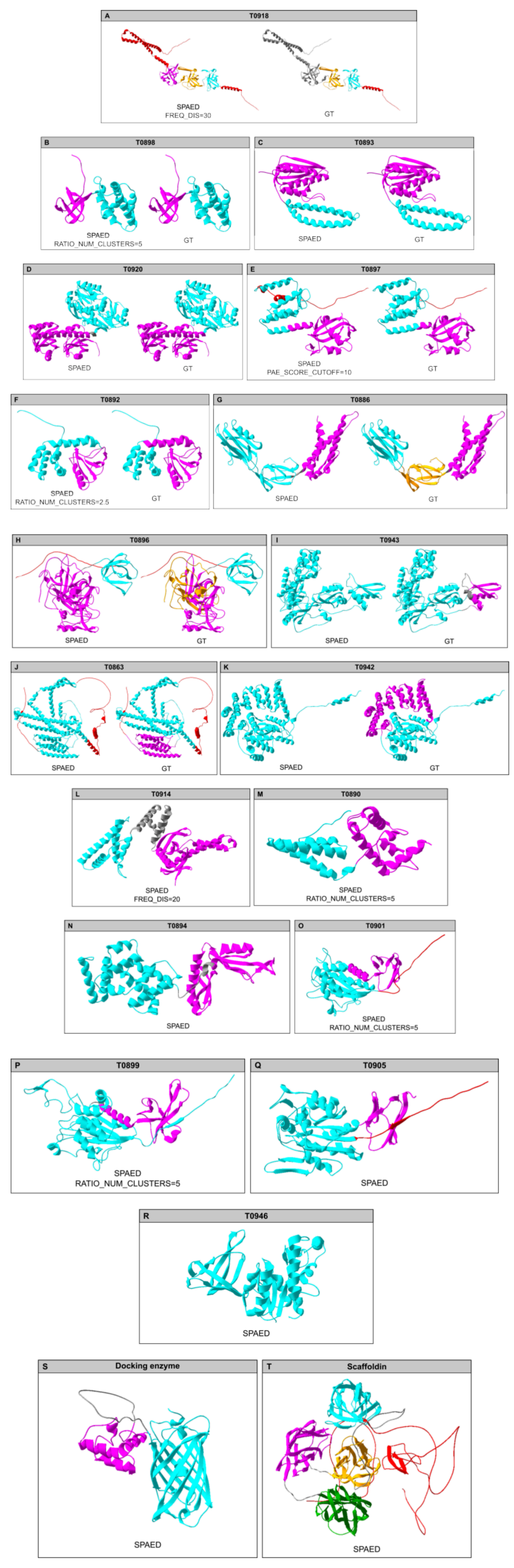
